# Shared local brain dynamics in pediatric and adult NREM parasomnias

**DOI:** 10.1101/2025.09.16.676512

**Authors:** Julian Amacker, Marco Veneruso, Matteo Pereno, Simone Ulzega, Samuel Wherli, Sven Hirsch, Lino Nobili, Silvia Miano, Mauro Manconi, Anna Castelnovo

## Abstract

Disorders of arousal (DoA), a group of Non-Rapid Eye Movement parasomnias—including sleepwalking, night terrors, and confusional arousals—arise from incomplete awakenings during slow-wave sleep, yet their neural signatures remain poorly defined. Using high-density EEG and source-space spectral analysis in both children and adults, we mapped cortical dynamics in the seconds before and after DoA onset and compared them to physiological motor arousals. Across ages, DoA episodes emerged from a globally less activated cortical state, with pre-onset surges in delta and beta power peaking in premotor, orbitofrontal, and anterior cingulate cortices. After onset, episodes showed widespread beta enhancement and focal delta suppression in sensorimotor and parietal associative regions relative to stable slow-wave sleep, alongside sustained frontal delta and beta activity compared with physiological motor arousals. These age-invariant spatial patterns identify a stable neurophysiological signature of DoA, supporting the concept of local sleep–wake dissociation. Our findings provide a framework for mechanistic models and potential biomarkers to modulate, predict, and differentiate DoA from other nocturnal events.

## Introduction

Sleepwalking, sleep terrors, and confusional arousals are non-rapid eye movement (NREM) sleep parasomnias^1^, collectively known as Disorders of Arousal (DoA). These episodes arise from incomplete awakenings during slow-wave sleep (SWS)^2^ and are characterized by complex behaviors—such as sitting up, walking, or speaking—without full awareness and with reduced responsiveness to the environment. While often benign, DoA can cause sleep disruption, psychosocial distress^3, 4^, accidental injury, and, in rare cases, legal consequences^5–7^. Despite their prevalence—up to 13% in childhood and 2% in adults^8, 9^—their underlying neurophysiology remains incompletely defined, limiting diagnostic precision and therapeutic options^10^.

Sporadic single-photon emission computed tomography (SPECT)^11^ and intracerebral EEG (SEEG) recordings during epilepsy surgery evaluations^12–15^ have shown that DoA episodes often feature a mixture of slow-wave “sleep-like” activity and faster “wake-like” rhythms, consistent with local sleep–wake dissociation^16^. However, these techniques are invasive or impractical for repeated, systematic evaluation across multiple episodes. Scalp EEG^17, 18^ offers greater accessibility but lacks the spatial resolution to fully map local cortical dynamics. Nearly all scalp EEG studies to date have focused on adults^17–29^, leaving it unclear whether these features reflect universal mechanisms or age-specific processes. Furthermore, the extent to which DoA differ from physiological arousal processes—particularly in the seconds following movement onset——remains poorly understood, partly due to the difficulty of retaining clean EEG during movement-rich events. Only one study has examined the post-onset phase in adults^20^, and none in children.

Here, we address three key gaps in DoA neurophysiology. First, we provide the first fine-grained, source-space mapping of cortical oscillatory dynamics before and after DoA onset in adults, leveraging high-density EEG—a technique enabling 3D source modelling of cortical activity with spatial resolution comparable to PET^30^. Second, we test whether these spatiotemporal patterns are conserved across age, indicating core mechanisms rather than age-specific phenomena, or whether they are specific to more severe cases that persist into adulthood. Third, we determine how DoA differ from physiological motor arousals in both children and adults—before and after movement onset—to clarify whether DoA follow an exaggerated, delayed, or qualitatively distinct arousal trajectory, irrespective of age.

## Results

### Participants

High-quality hd-EEG recordings were obtained from 22 adults (mean age 30.3 ± 5.2 years, 12 females) and 19 children (mean age 10.9 ± 3.0 years, 6 females). After exclusion for sleep apnea or epileptiform activity, the final analyses included 20 adults and 17 children. Clinical and vPSG features are summarized in Supplementary Tables 1–4. None of the participants were taking psychotropic medications or drugs on the night of the study.

### Behavioral characterization of NREM parasomnia episodes and typical motor arousals

Of 157 SWS motor events in adults, 55 were excluded for lack of consensus among scorers, 34 were identified as parasomnia episodes and 68 as physiological motor arousals. From 154 SWS motor events in children, 26 were excluded for the same reason, leaving 50 DoA episodes and 78 physiological motor arousals.

Parasomnia behaviors most often consisted of brief confusional arousals with eye opening (observed in 91% of adults, in 82% of children)—accompanied by sitting up, staring or perplexed gaze, and exploratory head movements (Supplementary Movie 1). Sleep talking occurred occasionally (29% in children, 9% in adults), as did hallucinatory behaviors, such as imaginary conversations or pointing at nonexistent objects (12% in children, none in adults). Loud screaming was noted in (8%) of children’s events (Supplementary Movie 2) while a single pediatric episode involved an attempt to sleepwalk (Supplementary Movie 3). Physiological motor arousals typically involved turning over in bed, adjusting bedclothes, or scratching with eyes closed. See Supplementary Table 5 for other details.

### Spectral power dynamics from SWS to NREM parasomnia episodes in adults

In the 5 s preceding movement onset, source-level analysis revealed a robust delta power increase relative to stable SWS baseline (−3 to −2 min). This increase involved a broad frontal– midline network including prefrontal, cingulate, and orbitofrontal cortices—specifically the anterior and mid-cingulate cortex, inferior frontal gyrus, frontopolar areas, and subcallosal cortex. Temporal and occipital contributions were more limited and predominantly left-lateralized (Figure 1A, Table 1).

**Figure 1.**
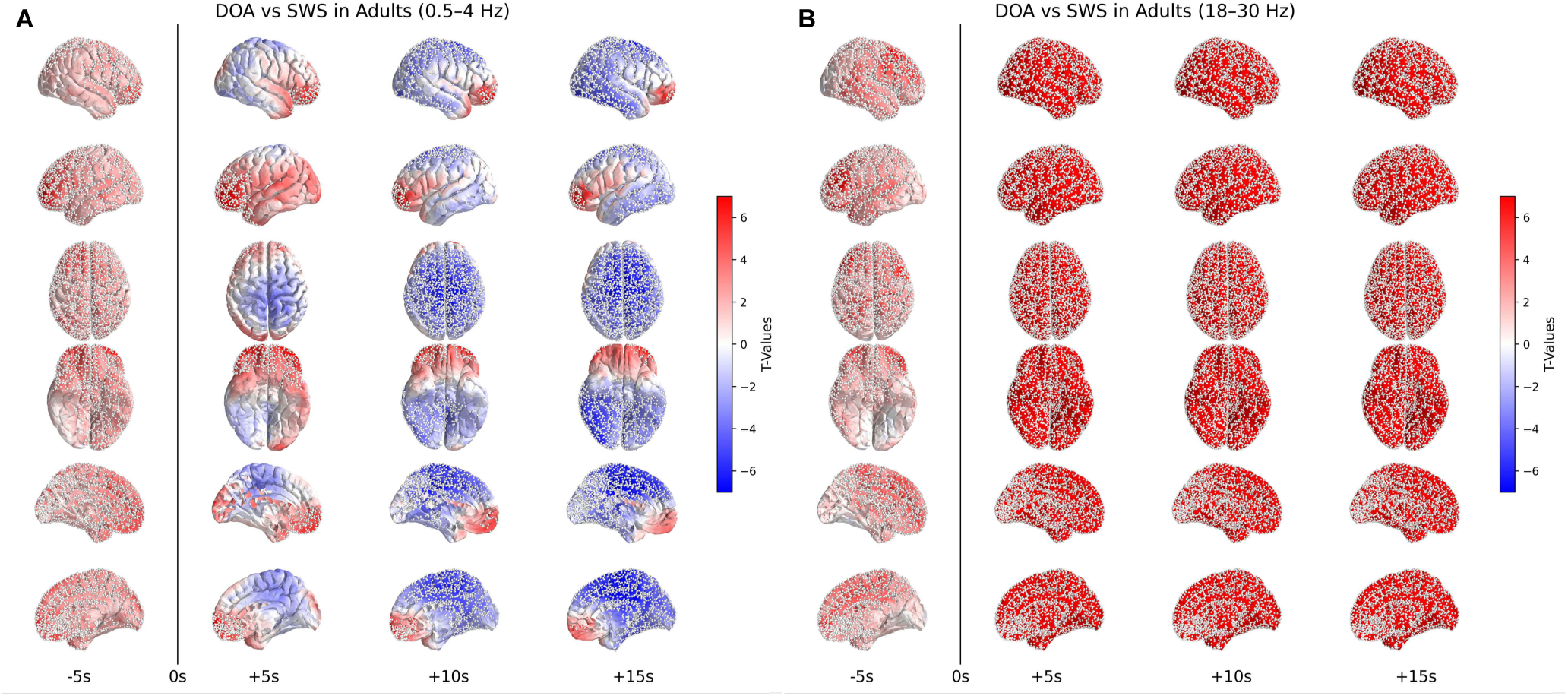
Spatiotemporal spectral dynamics around motor onset during DoA episodes in adults. *(A)* Source-level t-maps of delta power (0.5–4 Hz) show differences between DoA events and stable slow-wave sleep (SWS) baseline (−3 to −2 min), across four windows from −5 to +15 seconds relative to movement onset (0 s). Warmer colors (red) indicate increased power; cooler (blue) indicate decreased power during parasomnia. Notably, a widespread fronto-temporal/midline increase is seen at −5 s, followed by a central fronto-parietal suppression at +10 to +15 s. *(B)* Source-level t-maps of beta power (18–30 Hz) across the same windows show a symmetrical, widespread increase in activity prior and during DoA episodes. White dots indicate scouts with significant differences (pTFCE-corrected). All maps are thresholded by corrected t-values.

**Table 1.**
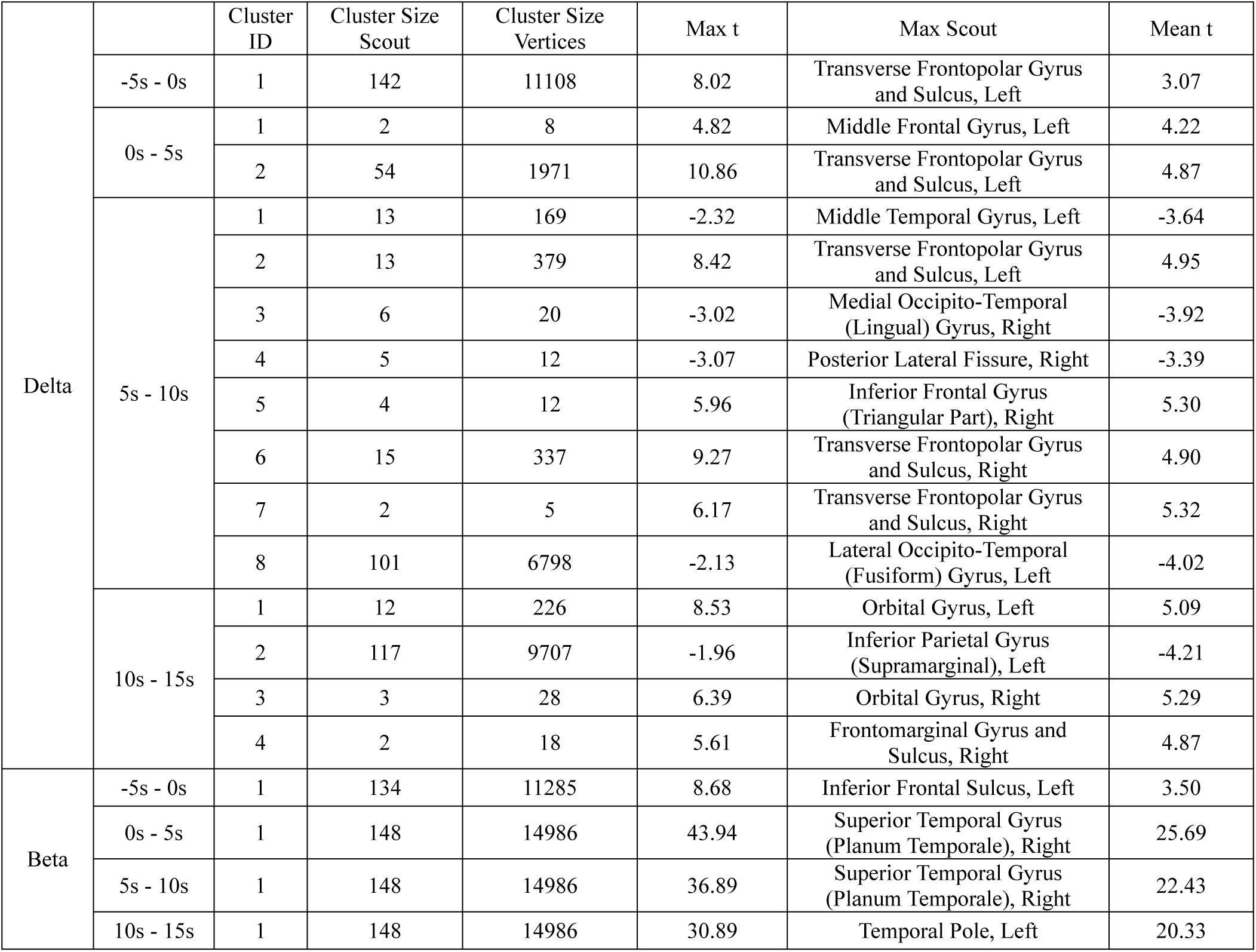
DoA episodes vs baseline in adults. Significant EEG clusters in Delta and Beta bands in the 20 seconds surrounding DoA episode onset (0s), compared to baseline slow wave activity recorded 2–3 minutes before the event. Results are organized in 5-second time bins. Legend: Time Windows: EEG data were analyzed in five consecutive 5-second bins: −5s to 0s, 0s to +5s, +5s to +10s, +10s to +15s relative to episode onset; Cluster Size (Scout): Number of anatomical scouts (brain regions) involved; Cluster Size (Vertices): Number of cortical vertices in the cluster.; Max Scout: Region with the peak statistical effect; Max/Mean t: Maximum and average t-statistics within each cluster.

Across the first 15 s after onset, frontal delta enhancement persisted—most prominently in frontopolar and anterior cingulate regions—including the transverse frontopolar gyri and sulci, fronto-marginal areas, orbitofrontal cortices, and anterior cingulate cortex, with the highest density of significant vertices. In parallel, a centro-parietal delta suppression emerged within 5 s, intensified over 5–15 s, and encompassed precuneus, superior parietal, and parieto-occipital cortices, precentral and frontomedial cortices (frontomarginal and paracentral regions), bilateral posterior and marginal cingulate cortex and occipitotemporal areas.

In the beta band, widespread bilateral power increases were already present before onset, involving the anterior and mid-cingulate cortex, precentral, postcentral, and paracentral gyri, as well as frontal areas, the temporopolar cortex, and superior temporal regions. The occipital pole and the inferior occipital and temporal surfaces (Figure 1B, Table 1) were largely spared. Beta activity remained significantly elevated throughout the 15 s post-onset window, with stable bilateral distribution.

### Spectral power spatial dynamics from SWS to NREM parasomnia episodes in children

In children, source-level analysis showed a robust, spatially organized delta increase relative to SWS with left-hemispheric dominance. As in adults, the right posterior lateral temporal cortex did not reach significance (Figure 2A, Table 2). In the first 5 s after onset, the delta build-up was broader than in adults, engaging frontal, temporal, occipital, cingulate and insular cortices, with particularly strong bilateral activity in orbitofrontal and frontopolar regions, anterior/mid-anterior cingulate, inferior/middle frontal gyri, insula and subcallosal areas. By contrast, a midline–sensorimotor cluster centered on pre-, post- and para-central gyri— appeared relatively spared immediately post-onset (negative but non-significant). This cluster became significantly suppressed between 5–15 s post-onset, with additional effects in the precuneus, mid/posterior cingulate, superior/middle frontal gyri bilaterally, and right inferior parietal cortex (angular and supramarginal; right > left).

**Figure 2.**
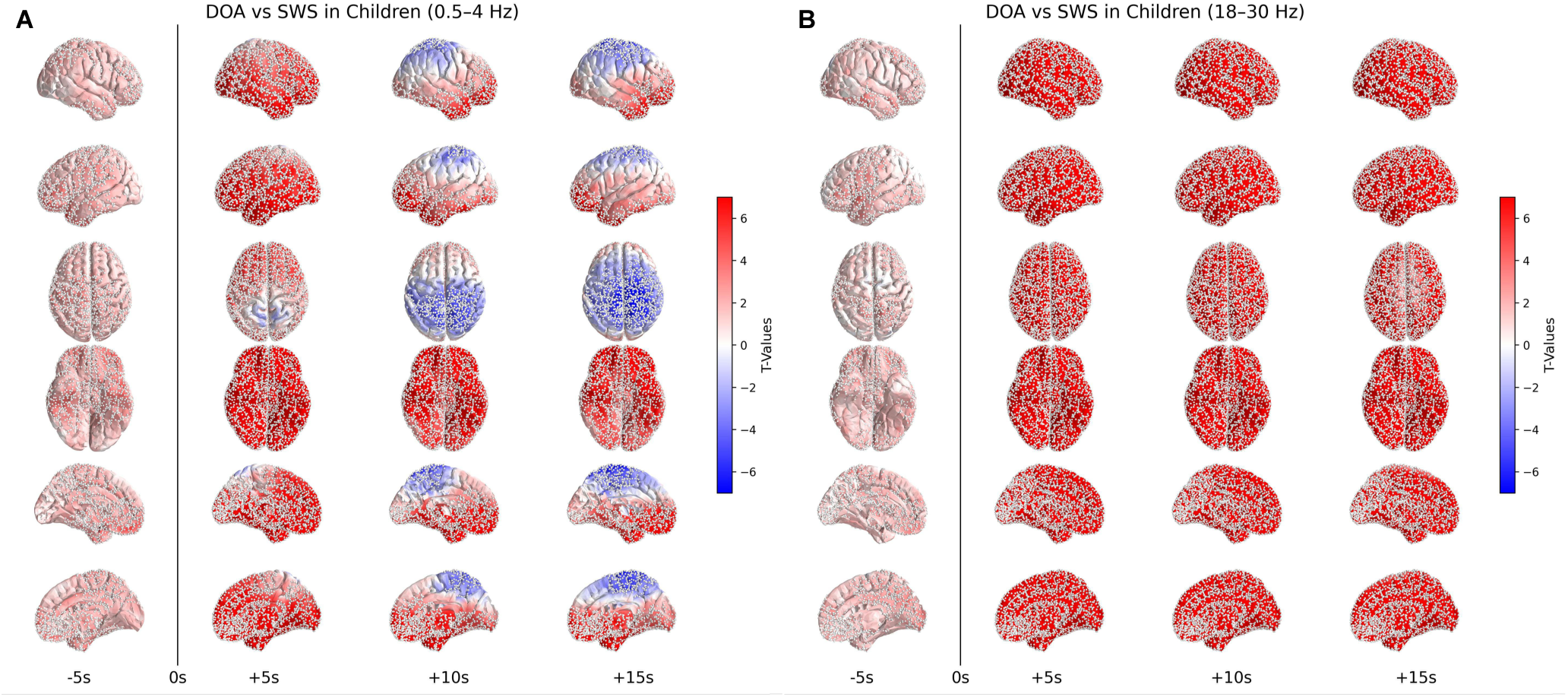
Spatiotemporal spectral dynamics around motor onset during DoA episodes in children. *(A)* Source-level t-maps of delta power (0.5–4 Hz) show differences between DoA events and stable slow-wave sleep (SWS) baseline (−3 to −2 min), across four windows from −5 to +15 seconds relative to movement onset (0 s). Warmer colors (red) indicate increased power; cooler (blue) indicate decreased power during parasomnia. In children, early widespread delta increase is observed at −5 s, followed by a fronto-parietal suppression from +10 s onward. Compared to adults (Figure 1A), the delta increase appears more diffuse and sustained, while the later suppression is more spatially confined. *(B)* Source-level t-maps of beta power (18–30 Hz) reveal sustained and bilateral increases in activity throughout the same windows, relative to SWS baseline. Compared to adults (Figure 1B), children show a similarly widespread and persistent beta enhancement across all time points. White dots indicate scouts with statistically significant differences (pTFCE-corrected). All maps are thresholded by corrected t-values.

**Table 2.**
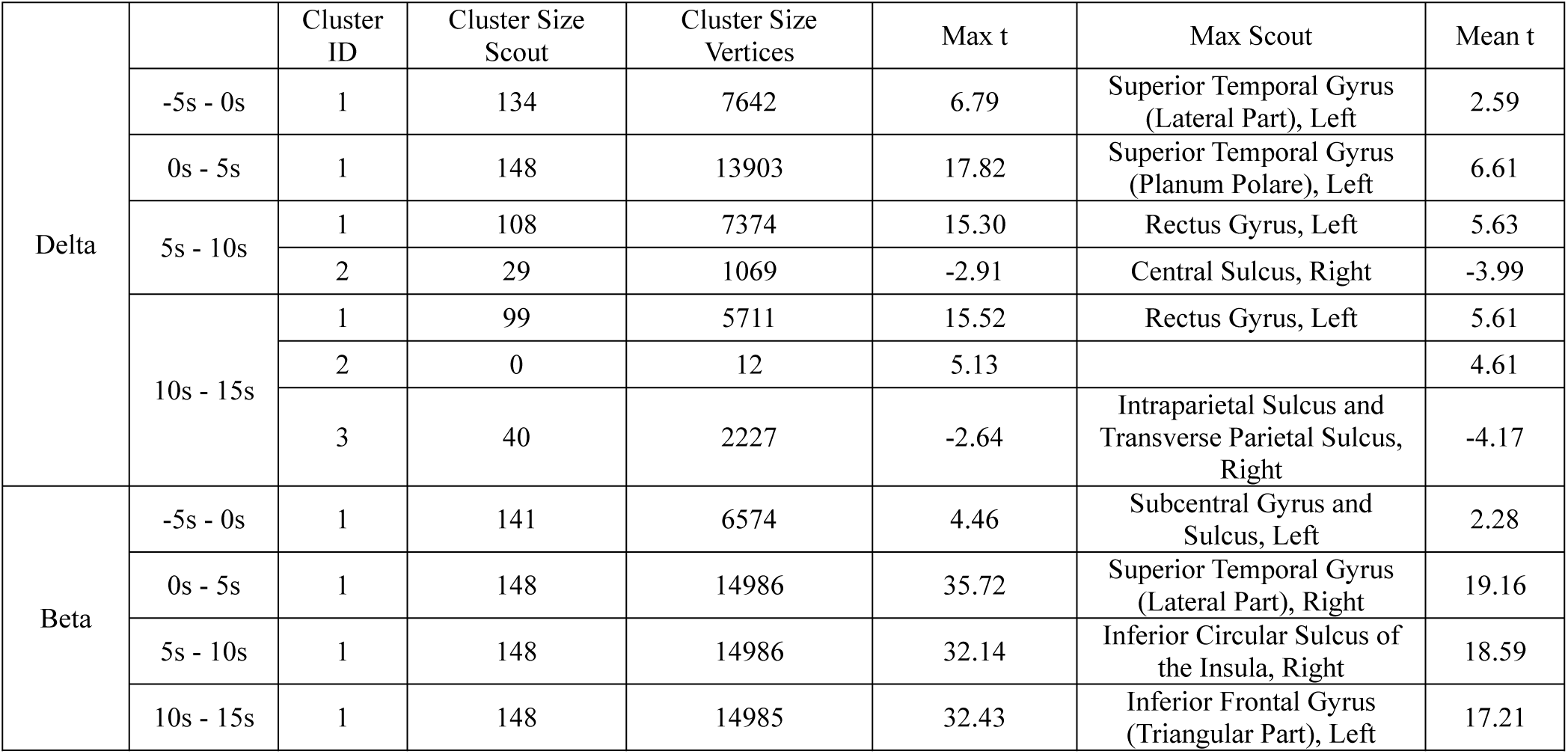
DoA episodes vs Baseline in children. Significant EEG clusters in Delta and Beta bands in the 20 seconds surrounding DoA episode onset (0s), compared to baseline slow wave activity recorded 2–3 minutes before the event. Results are organized in 5-second time bins. Legend: Time Windows: EEG data were analyzed in five consecutive 5-second bins: −5s to 0s, 0s to +5s, +5s to +10s, +10s to +15s relative to episode onset; Cluster Size (Scout): Number of anatomical scouts (brain regions) involved; Cluster Size (Vertices): Number of cortical vertices in the cluster.; Max Scout: Region with the peak statistical effect; Max/Mean t: Maximum and average t-statistics within each cluster

In the −5 to 0 s window, beta increases were less extensive than in adults, peaking in the left subcentral area and extending to inferior frontal and orbitofrontal regions, posterior–dorsal cingulate, insula, temporopolar cortex and part of the lateral temporal surface (left > right) (Figure 2B, Table 2). Post-onset, as in adults, beta activity was substantial and widespread throughout 0–15 s.

### Spectral power spatial differences surrounding NREM parasomnia episodes versus cortical motor arousals in children and adults

In both children and adults, delta power in the −5 to 0 s window was higher during DoA than physiological motor arousals, reaching statistical significance only in children, with the strongest effects over posterior cortices (Figure 3A,C; Tables 3–4). Beta power in the same window tended to be lower in DoA across similar regions, but differences were not significant after automatic cleaning (Figure 3B,D; Tables 3–4).

**Figure 3.**
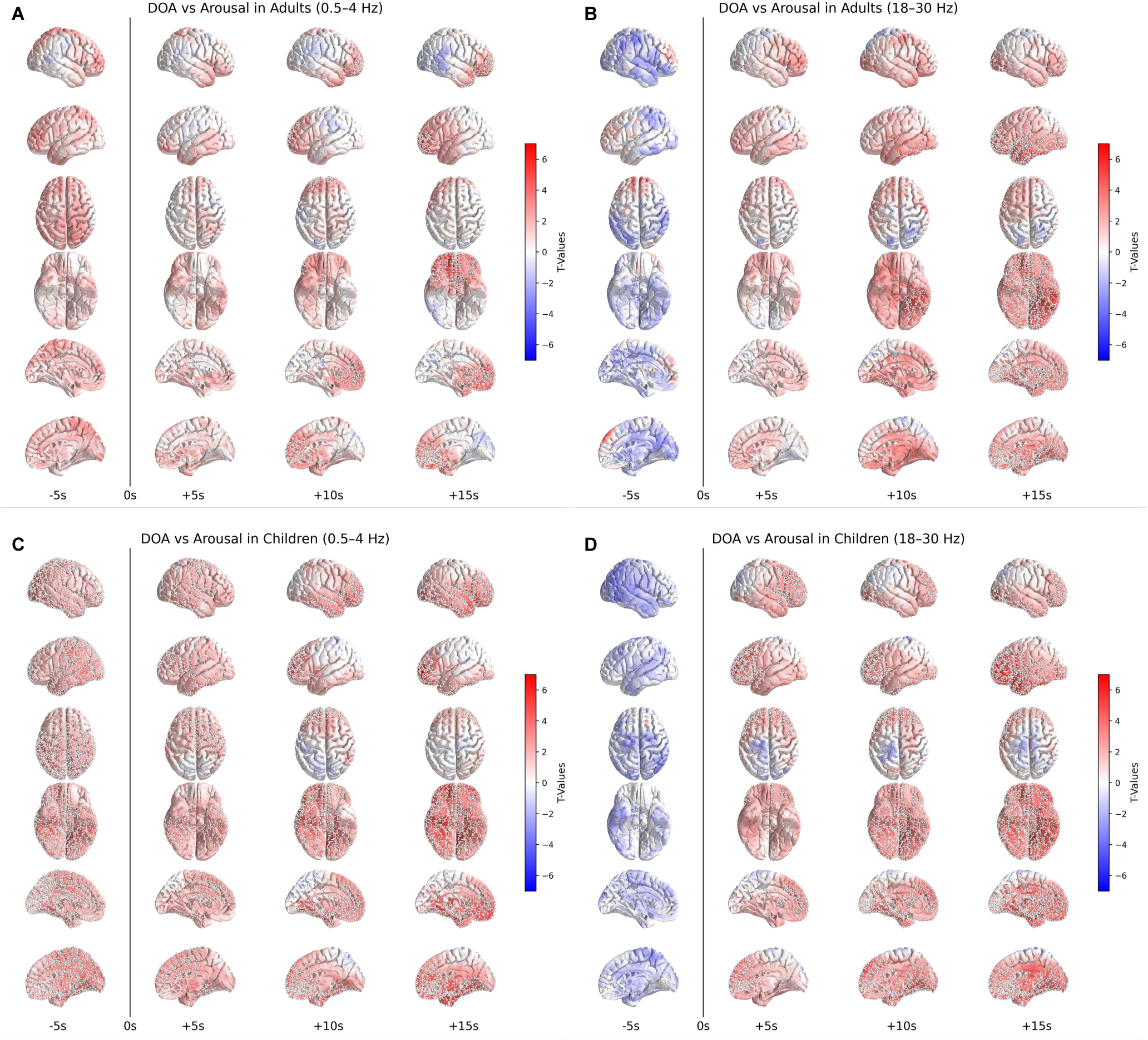
Comparison of spectral power between DoA events and typical motor arousals in adults and children. Source-level t-maps display spectral power differences between DoA episodes and typical motor arousals during the −5 to +15 second window relative to movement onset (0 s). Warmer colors (red) indicate higher power in parasomnia events compared to typical arousals; cooler colors (blue) indicate lower power. White dots mark vertices within scouts showing statistically significant differences (pTFCE-corrected t-values). *(A)* Delta power (0.5–4 Hz) in adults. Before movement onset, DoA events show higher delta activity, particularly in frontal and midline parietal regions. From 0 to +5 seconds, this difference becomes statistically significant over fronto-polar and fronto-basal surfaces. *(B)* Beta power (18–30 Hz) in adults. In the −5 to 0 second window, beta activity is lower during DoA events, especially over centro-posterior regions, but increases after motor onset, becoming significant between +5 and +10 seconds over ventral brain regions. *(C)* Delta power (0.5–4 Hz) in children. As in adults, DoA events in children are preceded by increased delta activity, which is widespread during the −5 to 0 second window and becomes confined to anterior brain regions post-onset—mainly the frontal cortex, anterior cingulate, and temporopolar areas. *(D)* Beta power (18–30 Hz) in children. Like adults, children show reduced beta activity during DoA events before onset and significantly higher beta during episodes, especially over ventral, frontal, and temporopolar brain regions.

**Table 3.**
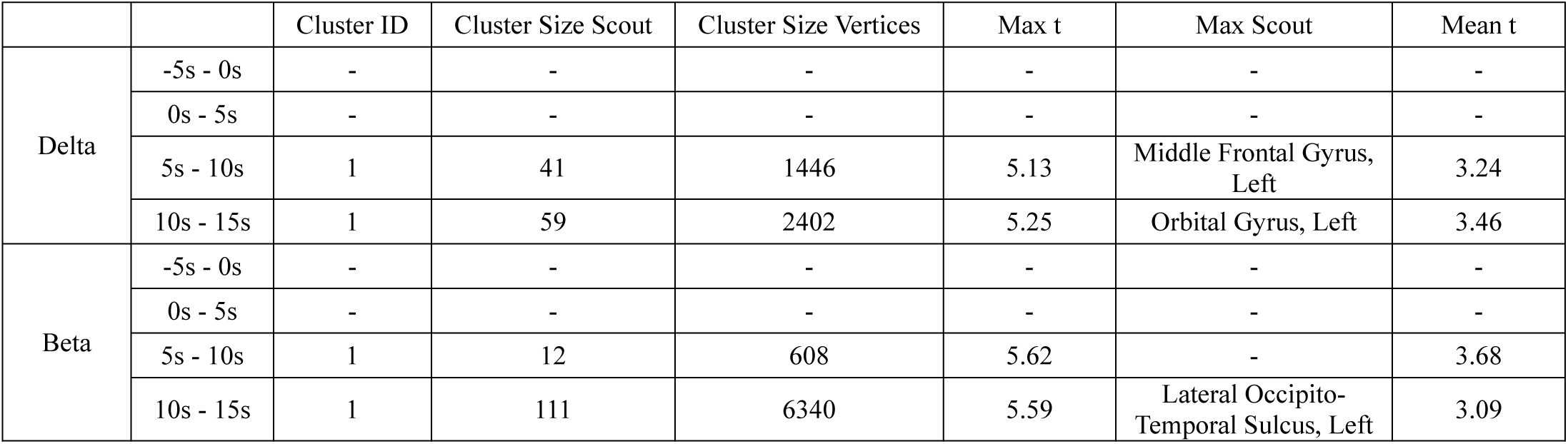
DoA episodes vs typical motor arousal in adults. Significant EEG clusters in Delta and Beta bands in the 20 seconds surrounding DoA episode onset (0s), compared to a similar EEG window surrounding physiological motor arousals. Results are organized in 5-second time bins. Legend: Time Windows: EEG data were analyzed in five consecutive 5-second bins: −5s to 0s, 0s to +5s, +5s to +10s, +10s to +15s relative to episode onset; Cluster Size (Scout): Number of anatomical scouts (brain regions) involved; Cluster Size (Vertices): Number of cortical vertices in the cluster; Max Scout: Region with the peak statistical effect; Max/Mean t: Maximum and average t-statistics within each cluster.

Following onset, delta power remained higher in DoA. In children, effects were widespread, with positive clusters in dorsal frontal, anterior cingulate, and ventral cortical areas, sparing a centro-posterior zone described above. Adults showed a similar but more anteriorly focused pattern, strongest between 5–15 s post-onset (Figure 3A,C; Tables 3-4).

**Table 4.**
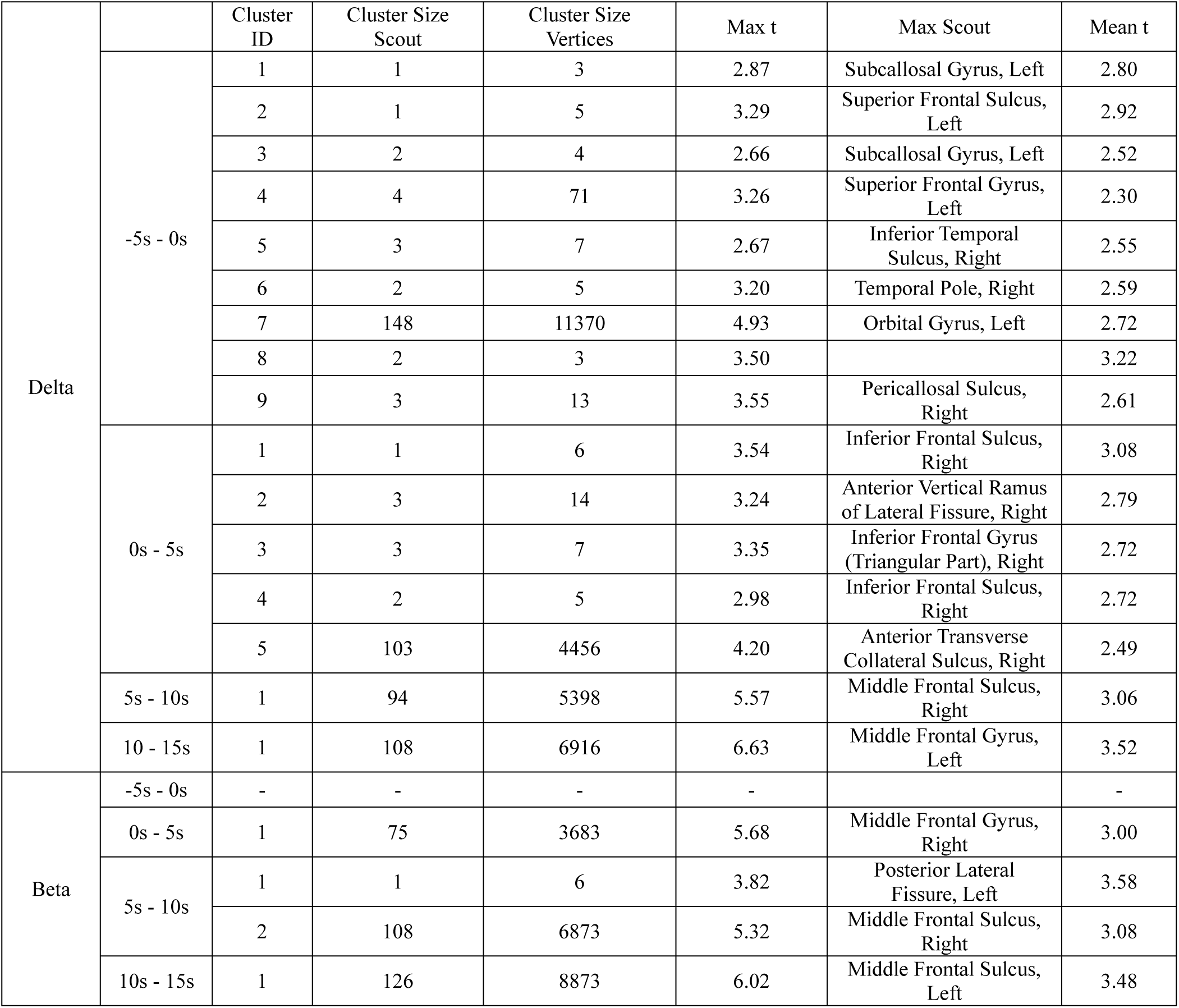
DoA episodes vs typical motor arousal in children. Significant EEG clusters in Delta and Beta bands in the 20 seconds surrounding DoA episode onset (0s), compared to a similar EEG window surrounding physiological motor arousals. Results are organized in 5-second time bins. Legend: Time Windows: EEG data were analyzed in five consecutive 5-second bins: −5s to 0s, 0s to +5s, +5s to +10s, +10s to +15s relative to episode onset; Cluster Size (Scout): Number of anatomical scouts (brain regions) involved; Cluster Size (Vertices): Number of cortical vertices in the cluster; Max Scout: Region with the peak statistical effect; Max/Mean t: Maximum and average t-statistics within each cluster.

Post-onset beta power was significantly higher in DoA than physiological motor arousals, with an early diffuse rise followed by more focal anterior and ventral enhancements. This effect was robust in children; in adults, significant clusters were fewer and appeared mainly between 5– 15 s post-onset (Figure 3B,D; Tables 3-4).

## Discussion

Using high-density EEG, we mapped cortical oscillatory dynamics in DoA with unprecedented spatial detail, resolving activity changes in 5-second windows before and after movement onset. By studying both children and adults and directly contrasting DoA with physiological motor arousals, we establish a quantitative, anatomically precise framework for mechanistic interpretation. This approach uncovered three key advances: robust regional brain activity changes from SWS during DoA, clear differentiation from physiological motor arousals, and consistent pre- and post-onset patterns across age groups.

In the seconds immediately preceding a DoA episode, we observed a pronounced delta increase in both children and adults, peaking in post-central, cingulate and frontal areas, likely reflecting arousal-related transitions in brain state. Concurrent beta increases mainly engaged motor-associative networks, consistent with action readiness or early motor planning. These changes expand on earlier low-resolution EEG studies^18, 22–27, 29^ by providing a detailed topographical characterization and revealing qualitative similarities with physiological arousals^31–35^.

In the seconds after a DoA episode onset, both children and adults showed widespread beta enhancement and persistent delta increases—most prominent in the anterior prefrontal cortex—alongside focal delta suppression over centro-parietal regions. This suppression was strongest in areas implicated in motor planning and executive control^36^, visuospatial integration^37^, self-referential processing, multimodal sensory integration^38, 39^, the default-mode and attentional networks^40, 41^ and conscious experiences during both dreaming and NREM parasomnias^21, 42^. The described cluster of delta suppression closely resembles that reported by Cataldi et al. ^21^ when contrasting episodes with and without conscious experiences, consistent with their finding that the former are more frequent. The weaker centro-parietal suppression in children supports, albeit on limited evidence^43, 44^, he view that mental activity is less common in pediatric episodes. The concurrent low- and high-frequency increases in distinct, though partially overlapping, cortical regions^12^, mirror stereo-EEG finding^45^, where wake-like motor cortex activation was preceded by frontal delta bursts during both subtle and overt movements^46^. Together, these findings suggest a functional dissociation between motor-control and executive-awareness networks—a hallmark of DoA that enables complex motor acts with diminished environmental responsiveness.

When directly compared with physiological motor arousals, DoA arose from a more “sleep-like” background, with globally higher delta and lower beta power—a pattern consistent with previous adult findings^20, 47^, and even more pronounced in children. Post-onset, differences became more spatially localized, with robust low- and high-frequency activation in frontotemporal-polar and limbic regions, particularly the anterior cingulate. These findings suggest that DoA involve a qualitative disruption in integrating sensorimotor and higher-order executive networks.

Finally, our results reveal that the temporal and spatial pre- and post-onset EEG patterns of DoA are strikingly consistent across age groups. This cross-age continuity supports the idea that DoA share a stable core neurophysiological mechanism, rather than representing age-specific or developmentally distinct phenomena. The persistence of episodes into adulthood may reflect a greater genetic predisposition or, as shown by several studies, the continued presence or emergence of predisposing or triggering factors, such as stress or psychopathology^3^.

Our regional delta dynamics partially diverge from Cataldi et al.^20^, who found a global delta reduction during adult DoA episodes and less consistent, less localized post-onset effects. This discrepancy likely reflects methodological differences: we analyzed only baseline-night arousals, applied more conservative artifact rejection (manual channel/epoch cleaning plus milder automatic procedures), and performed source-space analyses on normalized data using mixed-effects modeling. The consistency of our patterns across children and adults, and their close replication of previous findings from a pediatric case report^48^ and stereo-EEG studies (Supplementary Table 6), supports the robustness and reproducibility of these spatiotemporal dynamics.

Taken together, our findings suggest that, while local sleep–wake dissociation is an adaptive NREM feature—allowing brief motor adjustments without full awakening—it persists abnormally during DoA. Before motor onset, differences between DoA and physiological motor arousals are primarily quantitative and global rather than qualitative and regional, supporting the view that DoA often occur during predisposing time windows when cortical activation is insufficient for full wakefulness. Depending on cortical state (e.g., a “sleepier” background shaped by NREM depth or infra-slow oscillations), the strength and pattern of subcortical arousal activation, and individual predisposition (e.g., genetic load), arousal may resolve spontaneously, transition to full wakefulness, or escalate into a DoA episode. What distinguishes DoA is not the initiation of arousal, but the failure to engage key regions—such as the prefrontal cortex and anterior cingulate—needed for full awakening and behavioral regulation. Rather than a deficit in arousal intensity, DoA may represent a breakdown in arousal integration across cortical subsystems, producing partial awakenings in which sensorimotor circuits reactivate while higher-order networks remain offline.

We speculate that this cortical-level dissociation may arise from altered activity in the wake-promoting noradrenergic system^49–51^, whose modular organization could differentially support motor and prefrontal circuits^51, 52^. In line with ultra–high-field fMRI studies on sleep arousal ^33^, DoA may reflect an incomplete progression through the thalamic arousal sequence: early hubs, such as the VIM—active during DoA^15^—are engaged, supporting core arousal^53, 54^, while later nuclei, including the pulvinar—partially inactive in DoA^12^—remain offline. These later nuclei mediate integrative motor, cognitive, and sensory functions^55^, and their recruitment depends on synchronized activation of associative cortical areas, typically occurring only after diffuse arousal nuclei are engaged^56–58^. In DoA, the process may stall before this stage, due to an impaired frontal activation, preventing full awakening of late thalamic hubs and of higher-order cortical networks.

Limitations include grouping multiple parasomnia subtypes—most events were confusional arousals—and using template MRIs for source modelling, which may slightly reduce spatial precision. Although subgroup analyses were not feasible, future studies should clarify whether cortical dynamics differ among confusional arousals, sleepwalking, and night terrors to strengthen generalizability. Moreover, as all participants were recruited from a specialized sleep medicine center, the sample may be biased toward more severe or treatment-seeking cases, potentially limiting applicability to milder community presentations. However, the high reproducibility across ages and convergence with invasive studies support the robustness of our patterns.

Future studies with larger samples, individual MRI data, and combined behavioral, neurophysiological, and phenomenological measures are needed to confirm these findings and address remaining questions on the neural dynamics of physiological and pathological DoA. In particular, integrating hd-EEG data with subjective reports in children may help disentangle arousal-related circuits from networks sustaining consciousness.

To conclude, our source mapping analysis is consistent with, yet expands upon, previous literature, providing a detailed neurophysiological framework for investigating DoA mechanisms. The growing recognition of the modularity of arousal systems—at both modulatory neurotransmitter and thalamocortical levels—could guide the development of new, testable hypotheses based on the precise topographical patterns identified here, advancing the understanding of DoA pathophysiology and informing novel pharmacotherapeutic approaches. Furthermore, the identification of EEG changes that precede movement onset may enable future efforts to predict or modulate episodes through spectral EEG dynamics, support early intervention or preventive strategies in high-risk individuals, and improve differential diagnosis with sleep-related hypermotor epilepsy (SHE) when only minor episodes are captured in the laboratory.

## Methods

This observational, intra-group, monocentric study was conducted at the Sleep Medicine Unit of the Neurocenter of Southern Switzerland at the Civic Hospital of Lugano between July 2018 and April 2023. All study procedures were reviewed and approved by the Local Ethics Committee (2017-01788 CE 3282) and conducted in accordance with the Declaration of Helsinki.

### Participants

Patients were recruited either from outpatients attending our sleep medicine center (during routine visits or retrospectively through the electronic hospital system) or from the general population via word of mouth. Inclusion criteria comprised a clinical diagnosis of DoA based on ICSD-3 criteria^2^ and an age range of 5–17 years (children’s group) or 18–45 years (adult group). Exclusion criteria included major psychiatric or neurological conditions (e.g., epilepsy), use of psychotropic medications, and an apnea-hypopnea index (AHI) >5 events/hour for children or >15 events/hour for adults. Following an in-person or telephone screening to assess eligibility, all patients—and, when available, their parents or bedpartners— underwent a comprehensive clinical evaluation by a physician board-certified in sleep medicine. Relevant clinical data, including age, sex, family history of parasomnia, age of onset, and episode frequency at the time of observation, were collected. Informed consent was obtained from all participants prior to study initiation.

### Experimental procedure

All enrolled patients underwent at least one nocturnal video-polysomnography (PSG) recording in our sleep laboratory (∼09:00 pm to ∼07:00 am). Non-EEG channels were recorded using an Embla N7000 PSG System and included video, electro-oculogram (EOG), electromyogram (EMG) of the submentalis muscle, the right and left tibialis anterior muscles and extensor digitorum communis, electrocardiogram, oral and nasal airflow thermistors, nasal pressure cannula, and wearable piezo-electric bands for thoracic and abdominal movements. EEG was recorded using a high-density EEG system (256 channels; Electrical Geodesics Inc., Eugene, OR, vertex referencing, sampling rate of 500 Hz).

Technicians intervened in DoA episodes only if the episode lasted more than 20-30 seconds. In such cases, the experimenter called the patient’s name and asked simple questions, such as “Where are we?” and “Can you name this object?”.

### Sleep scoring

Sleep staging and the scoring of associated sleep events was performed by a board-certified sleep physician (AC) according to standard AASM criteria^59^ using EMBLA-RemLogic software. Sleep staging was based on 30-second epochs for 6 standard EEG derivations extracted from the high-density EEG system, with bipolar re-referencing (F3/M2, F4/M1, C3/M2, C4/M1, O1/M2, O2/M1), and synchronized with the other PSG channels.

### Scoring of parasomnia episodes and of normal arousals

DoA episodes were defined as sudden behaviors emerging against a sleep EEG background and characterized by eye opening and/or vocalizations, movements of the head or limbs, trunk flexion, exploratory actions, or abrupt expressions of fear. Typical motor arousals were identified as EEG arousals associated with physiological movements, such as changing body position, adjusting posture, scratching, stretching, and rearranging clothing or blankets. To enhance the accuracy and consistency of evaluating behavioral episodes, three raters certified in sleep medicine (AC, SM, LN for children; AC, MV and LN for adults) independently viewed all the video-EEG recordings and classified the behavior as either a parasomnia episode or a typical motor arousal. Discordant cases were resolved by discussion. Events for which the three experts could not reach a consensus were excluded from the analysis.

The beginning of a motor event (either a parasomnia episode or a typical motor arousal) was defined as the onset of movement detected on one of the five EMG channels available (submental—chin, bilateral arms, bilateral legs), whichever occurred first. The end of a motor event was determined by the definite cessation of movement, the appearance of a clear wake EEG pattern, or the patient’s ability to correctly answer two consecutive questions in cases where the technician intervened. Events with brief interruptions of motor activity, lasting fewer than 5 seconds and followed by the resumption of movement, were considered a single episode.

### EEG data extraction

EEG data analysis was performed in MATLAB (The MathWorks Inc., Natick, MA, USA) using the EEGLAB v13^60^.

Initially, 15-minute EEG segments were extracted, comprising 10 minutes of sleep EEG preceding motor onset and 5 minutes following motor onset. These segments were then band-pass filtered between 0.5 and 45 Hz. To avoid edge distortion introduced by filtering, shorter 5-minute segments were subsequently retained, spanning from 3 minutes before to 2 minutes after motor onset. This step was taken to avoid edge distortion caused by filtering.

### EEG artifact removal procedure

Channels affected by artifacts (either for the entire segment or for 5-second sub-epochs) were visually inspected and labeled as bad in Brainstorm^61^. Artifact Subspace Reconstruction (ASR) was then applied, after excluding identified bad channels. ASR adaptively filters out artifacts by modeling clean EEG segments—automatically selected based on deviations from the power distribution in each channel—and rejecting outliers in the PCA subspace. A moving window of 768 ms (1.5 × the number of channels) and a 35 standard deviation cutoff for rejection were used to avoid excluding sleep features, such as slow waves persisting during the arousal/episode. This threshold is consistent with the default value of the Dusk2Dawn automatic cleaning procedure^62^.

To remove ECG and muscle artifacts, wavelet-enhanced ICA (wICA) was applied using a customized version of the EEGLAB “RELAX” plugin^63^. ICLabel classification was adapted for hd-EEG by labeling components as artifactual if classified as ‘bad’ (excluding ‘brain’ and ‘other’) with >70% probability. This selectively removed noise while preserving neural signals. The same wICA, ASR, and ICA procedures were used for both parasomnia episodes and typical motor arousals.

### Power Spectral Density decomposition

Power spectral density (PSD) was computed at the source level using Welch’s modified periodogram method, with 5-second Hamming windows and 50% overlap. The analysis focused on the average PSD magnitude within the delta (0.5–4 Hz) and beta (18–30 Hz) frequency bands. At the scalp level, bad channels were interpolated using spherical spline. For source-level analysis, only normalized maps obtained via sLORETA-based minimum norm estimates were considered.

### Source modeling

Source modeling was performed with Brainstorm using an age-appropriate template^64^, segmented using SPM12/CAT12 Matlab toolbox^65^. A symmetric Boundary Element Method (BEM) volume conduction model of the head having three realistic layers (scalp, inner skull, outer skull)^29^ and a standard co-registered set of electrode positions were used to construct the forward model. The inverse matrix was computed using the sLORETA Minimum Norm^66^ with sources constrained to be perpendicular to the cortical surface and retaining only diagonal elements of the noise covariance matrix.

### Statistical analyses

Statistical comparisons between DoA episodes and baseline sleep were performed using a linear mixed-effects model (LME) for each electrode or voxel and epoch, using the lmeEEG Matlab routine^67^. This method reduces computational load by isolating marginal EEG data for standard regression.

Two primary comparisons were conducted: DoA/baseline and DoA/motor movement. Dependent variables included EEG-derived measures such as scalp or source PSD. Episode onset (from sleep onset) and age were included as fixed effects, whereas adding sex yielded no substantive improvement in ΔAIC and ΔBIC. We therefore adopted the more parsimonious model excluding sex. Subjects were treated as a random factor to account for the fact that participants contributed multiple episodes. To create the null distribution, condition labels were shuffled within subjects: 2000 iterations for the DoA/baseline contrast and 3000 iterations for the DoA/motor contrast - the latter increased because p-values lay closer to the significance threshold and thus required a more robust estimate. For the DoA/motor model, lmeEEG was further adapted to implement the Freedman–Lane permutation scheme, ensuring valid permutations even for participants who contributed only one of the two conditions (DoA or motor)^68^.

Probabilistic Threshold-Free Cluster Enhancement (pTFCE)^69^ was applied to correct for multiple comparisons using empirically derived parameters for height and extent (E = 2, H = 0.66 for PSD).

## Data availability

The datasets generated during and/or analyzed during the current study are available from the corresponding author on reasonable request. Only a subset of parasomnia episode videos is included in the supplementary materials. Additional videos can be accessed upon direct request to the corresponding author, in accordance with patient consent.

## Code availability

Templates of the custom-made codes in Matlab are available at https://github.com/julama/hdEEG_DOA.

## Acknowledgements

This work was supported by the EOC Young Researchers Grant 2023 awarded to A.C. The authors thank the patients and their families for participation in the study, and the entire EOC neurophysiology technical team for help with data acquisition.

## Author contributions

Conceptualization, A.C.; methodology, A.C.; software: J.A., A.C.; formal analysis, J.A., A.C., M.V., investigation, A.C., M.P.; resources: A.C., M.M.; data curation: M.V., J.A., A.C., L.N., S.M., writing (original draft), A.C., J.A., M.V.; writing (review and editing), all authors; visualization, J.A., M.V.; supervision: A.C., S.W., S.U.; project administration, A.C; and funding acquisition, A.C., S.H.

## Bibliography

1. Petit D, Touchette E, Tremblay RE, Boivin M, Montplaisir J. Dyssomnias and Parasomnias in Early Childhood. Pediatrics. 2007;119(5):e1016–e1025. doi:10.1542/peds.2006-2132

2. American Academy Of Sleep Medicine. International Classification of Sleep Disorders, Third Edition, Text Revision (ICSD-3-TR).; 2023.

3. Tomic T, Mombelli S, Oana S, et al. Psychopathology and NREM sleep parasomnias: A systematic review. Sleep Medicine Reviews. 2025;80:102043. doi:10.1016/j.smrv.2024.102043

4. Arnulf I, Zhang B, Uguccioni G, et al. A Scale for Assessing the Severity of Arousal Disorders. Sleep. 2014;37(1):127–136. doi:10.5665/sleep.3322

5. Siclari F, Khatami R, Urbaniok F, et al. Violence in sleep. Brain : a journal of neurology. 2010;133(Pt 12):3494–3509. doi:10.1093/brain/awq296

6. Castelnovo A, Schraemli M, Schenck CH, Manconi M. The parasomnia defense in sleep-related homicide: A systematic review and a critical analysis of the medical literature. Sleep medicine reviews. 2024;74. doi:10.1016/J.SMRV.2024.101898

7. Mahowald MW, Schenck CH, Goldner M, Bachelder V, Cramer-Bornemann M. Parasomnia pseudo-suicide. Journal of forensic sciences. 2003;48(5):1158–1162.

8. Petit D, Pennestri MH, Paquet J, et al. Childhood sleepwalking and sleep terrors: A longitudinal study of prevalence and familial aggregation. JAMA Pediatrics. 2015;169(7):653–658. doi:10.1001/jamapediatrics.2015.127

9. Stallman HM, Kohler M. Prevalence of Sleepwalking: A Systematic Review and Meta-Analysis. Published online 2016. doi:10.1371/journal.pone.0164769

10. Mainieri G, Loddo G, Provini F, Nobili L, Manconi M, Castelnovo A. Diagnosis and Management of NREM Sleep Parasomnias in Children and Adults. *Diagnostics (Basel*, Switzerland). 2023;13(7). doi:10.3390/DIAGNOSTICS13071261

11. Bassetti C, Vella S, Donati F, Wielepp P, Weder B. SPECT during sleepwalking. Lancet. 2000;356(9228):484–485. doi:10.1016/S0140-6736(00)02561-7

12. Flamand M, Boudet S, Lopes R, et al. Confusional arousals during non-rapid eye movement sleep: Evidence from intracerebral recordings. Sleep. 2018;41(10):1–11. doi:10.1093/sleep/zsy139

13. Terzaghi M, Sartori I, Tassi L, et al. Evidence of dissociated arousal states during nrem parasomnia from an intracerebral neurophysiological study. Sleep. 2009;32(3):409–412. doi:10.1093/sleep/32.3.409

14. Terzaghi M, Sartori I, Tassi L, et al. Dissociated local arousal states underlying essential clinical features of non-rapid eye movement arousal parasomnia: An intracerebral stereo-electroencephalographic study. Journal of Sleep Research. 2012;21(5):502–506. doi:10.1111/j.1365-2869.2012.01003.x

15. Sarasso S, Pigorini A, Proserpio P, Gibbs SA, Massimini M, Nobili L. Fluid boundaries between wake and sleep: Experimental evidence from Stereo-EEG recordings. Archives Italiennes de Biologie. 2014;152(2-3):169–177. doi:10.12871/0002982920142311

16. Castelnovo A, Lopez R, Proserpio P, Nobili L, Dauvilliers Y. NREM sleep parasomnias as disorders of sleep-state dissociation. Nature reviews Neurology. 2018;14(8):470–481. doi:10.1038/s41582-018-0030-y

17. Zadra A, Pilon M, Joncas S, Rompré S, Montplaisir J. Analysis of postarousal EEG activity during somnambulistic episodes. Journal of Sleep Research. 2004;13(3):279–284. doi:10.1111/j.1365-2869.2004.00404.x

18. Schenck CH, Pareja JA, Patterson AL, Mahowald MW. Analysis of polysomnographic events surrounding 252 slow-wave sleep arousals in thirty-eight adults with injurious sleepwalking and sleep terrors. Journal of Clinical Neurophysiology. Published online 1998. doi:10.1097/00004691-199803000-00010

19. Ratti PL, Amato N, David O, Manconi M. A high-density polysomnographic picture of disorders of arousal. Sleep. 2018;41(11). doi:10.1093/sleep/zsy162

20. Cataldi J, Stephan AM, Marchi NA, Haba-Rubio J, Siclari F. Abnormal timing of slow wave synchronization processes in non-rapid eye movement sleep parasomnias. Sleep. Published online May 2022. doi:10.1093/SLEEP/ZSAC111

21. Cataldi J, Stephan AM, Haba-Rubio J, Siclari F. Shared EEG correlates between non-REM parasomnia experiences and dreams. Nature communications. 2024;15(1). doi:10.1038/S41467-024-48337-7

22. Zadra A, Nielsen T. Topographical EEG mapping in a case of recurrent sleep terrors. Dreaming. Published online 1998. doi:10.1023/B:DREM.0000005897.62698.1b

23. Espa F, Ondze B, Deglise P, Billiard M, Besset A. Sleep architecture, slow wave activity, and sleep spindles in adult patients with sleepwalking and sleep terrors. Clinical Neurophysiology. 2000;111(5):929–939. doi:10.1016/S1388-2457(00)00249-2

24. Guilleminault C, Poyares D, Abat F, Palombini L. Sleep and wakefulness in somnambulism: A spectral analysis study. Journal of Psychosomatic Research. Published online 2001. doi:10.1016/S0022-3999(01)00187-8

25. Jaar O, Pilon M, Carrier J, Montplaisir J, Zadra A. Analysis of slow-wave activity and slow-wave oscillations prior to somnambulism. Sleep. 2010;33(11):1511–1516. doi:10.1093/sleep/33.11.1511

26. Perrault R, Carrier J, Desautels A, Montplaisir J, Zadra A. Electroencephalographic slow waves prior to sleepwalking episodes. Sleep Medicine. 2014;15(12):1468–1472. doi:10.1016/j.sleep.2014.07.020

27. Januszko P, Niemcewicz S, Gajda T, et al. Sleepwalking episodes are preceded by arousal-related activation in the cingulate motor area: EEG current density imaging. Clinical Neurophysiology. 2016;127(1):530–536. doi:10.1016/j.clinph.2015.01.014

28. Desjardins MÈ, Carrier J, Lina JM, et al. EEG functional connectivity prior to sleepwalking: Evidence of interplay between sleep and wakefulness. Sleep. 2017;40(4):1–8. doi:10.1093/sleep/zsx024

29. Mainieri G, Loddo G, Castelnovo A, et al. EEG Activation Does Not Differ in Simple and Complex Episodes of Disorders of Arousal: A Spectral Analysis Study. NSS. 2022;Volume 14:1097–1111. doi:10.2147/nss.s360120

30. Hedrich T, Pellegrino G, Kobayashi E, Lina JM, Grova C. Comparison of the spatial resolution of source imaging techniques in high-density EEG and MEG. NeuroImage. 2017;157:531–544. doi:10.1016/j.neuroimage.2017.06.022

31. Sforza E, Jouny C, Ibanez V. Cardiac activation during arousal in humans: Further evidence for hierarchy in the arousal response. Clinical Neurophysiology. Published online 2000. doi:10.1016/S1388-2457(00)00363-1

32. Peter-Derex L, Magnin M, Bastuji H. Heterogeneity of arousals in human sleep: A stereo-electroencephalographic study. NeuroImage. 2015;123:229–244. doi:10.1016/j.neuroimage.2015.07.057

33. Setzer B, Fultz NE, Gomez DEP, et al. A temporal sequence of thalamic activity unfolds at transitions in behavioral arousal state. Nat Commun. 2022;13(1). doi:10.1038/s41467-022-33010-8

34. Stephan AM, Cataldi J, Singh Virk A, Siclari F. Cortical activity upon awakening from sleep reveals consistent spatio-temporal gradients across sleep stages in human EEG. Current Biology. Published online July 2025. doi:10.1016/j.cub.2025.06.064

35. Massimini M, Ferrarelli F, Esser SK, et al. Triggering sleep slow waves by transcranial magnetic stimulation. Proc Natl Acad Sci USA. 2007;104(20):8496–8501. doi:10.1073/pnas.0702495104

36. Battaglia-Mayer A, Caminiti R. Corticocortical Systems Underlying High-Order Motor Control. J Neurosci. 2019;39(23):4404–4421. doi:10.1523/jneurosci.2094-18.2019

37. Xu Y, Chun M. Representing Visual Objects, Attention, and Load in Human Occipito-temporal and Posterior Parietal Cortices. Journal of Cognitive Neuroscience. Published online June 20, 2025:1–24. doi:10.1162/jocn.a.64

38. Sulpizio V, Fattori P, Pitzalis S, Galletti C. Functional organization of the caudal part of the human superior parietal lobule. Neuroscience & Biobehavioral Reviews. 2023;153:105357. doi:10.1016/j.neubiorev.2023.105357

39. Pflugshaupt T, Nösberger M, Gutbrod K, Weber KP, Linnebank M, Brugger P. Bottom-up Visual Integration in the Medial Parietal Lobe. Cereb Cortex. 2016;26(3):943–949. doi:10.1093/cercor/bhu256

40. Leech R, Sharp DJ. The role of the posterior cingulate cortex in cognition and disease. Brain. 2014;137(1):12–32. doi:10.1093/brain/awt162

41. Leech R, Kamourieh S, Beckmann CF, Sharp DJ. Fractionating the Default Mode Network: Distinct Contributions of the Ventral and Dorsal Posterior Cingulate Cortex to Cognitive Control. J Neurosci. 2011;31(9):3217–3224. doi:10.1523/jneurosci.5626-10.2011

42. Siclari F, Baird B, Perogamvros L, et al. The neural correlates of dreaming. Nature Neuroscience. 2017;20(6):872–878. doi:10.1038/nn.4545

43. Castelnovo A, Loddo G, Provini F, Miano S, Manconi M. Mental activity during episodes of sleepwalking, night terrors or confusional arousals: Differences between children and adults. Nature and Science of Sleep. 2021;13:829–840. doi:10.2147/NSS.S309868

44. Castelnovo A, Siclari F, Spaggiari S, et al. Conscious experiences during non-rapid eye movement sleep parasomnias. Neuroscience & Biobehavioral Reviews. 2024;167:105919. doi:10.1016/j.neubiorev.2024.105919

45. Ruby P, Eskinazi M, Bouet R, Rheims S, Peter-Derex L. Dynamics of hippocampus and orbitofrontal cortex activity during arousing reactions from sleep: An intracranial electroencephalographic study. Human Brain Mapping. 2021;42(16):5188–5203. doi:10.1002/hbm.25609

46. Nobili L, Ferrara M, Moroni F, et al. Dissociated wake-like and sleep-like electro-cortical activity during sleep. NeuroImage. 2011;58(2):612–619. doi:10.1016/J.NEUROIMAGE.2011.06.032

47. Castelnovo A, Mainieri G, Loddo G, et al. Spectral dynamics prior to motor events differ between NREM sleep parasomnias and healthy sleepers. SLEEP. 2025;48(3):zsae252. doi:10.1093/sleep/zsae252

48. Castelnovo A, Amacker J, Maiolo M, et al. High-density EEG power topography and connectivity during confusional arousal. Cortex. 2022;155:62–74. doi:10.1016/j.cortex.2022.05.021

49. Uematsu A, Tan BZ, Ycu EA, et al. Modular organization of the brainstem noradrenaline system coordinates opposing learning states. Nat Neurosci. 2017;20(11):1602–1611. doi:10.1038/nn.4642

50. Waterhouse BD, Chandler DJ. Heterogeneous organization and function of the central noradrenergic system. Brain Research. 2016;1641:v-x. doi:10.1016/j.brainres.2015.12.050

51. Chandler DJ, Gao WJ, Waterhouse BD. Heterogeneous organization of the locus coeruleus projections to prefrontal and motor cortices. Proc Natl Acad Sci USA. 2014;111(18):6816–6821. doi:10.1073/pnas.1320827111

52. Wagner-Altendorf TA, Fischer B, Roeper J. Axonal projection-specific differences in somatodendritic α2 autoreceptor function in locus coeruleus neurons. Eur J of Neuroscience. 2019;50(11):3772–3785. doi:10.1111/ejn.14553

53. Ilyas A, Pizarro D, Romeo AK, Riley KO, Pati S. The centromedian nucleus: Anatomy, physiology, and clinical implications. Journal of Clinical Neuroscience. 2019;63:1–7. doi:10.1016/j.jocn.2019.01.050

54. Vertes RP, Linley SB, Rojas AKP. Structural and functional organization of the midline and intralaminar nuclei of the thalamus. Front Behav Neurosci. 2022;16. doi:10.3389/fnbeh.2022.964644

55. Han J, Xie Q, Wu X, et al. The neural correlates of arousal: Ventral posterolateral nucleus-global transient co-activation. Cell Reports. 2024;43(1):113633. doi:10.1016/j.celrep.2023.113633

56. Vertes RP, Linley SB, Rojas AKP. Structural and functional organization of the midline and intralaminar nuclei of the thalamus. Front Behav Neurosci. 2022;16. doi:10.3389/fnbeh.2022.964644

57. Honjoh S, Sasai S, Schiereck SS, Nagai H, Tononi G, Cirelli C. Regulation of cortical activity and arousal by the matrix cells of the ventromedial thalamic nucleus. Nat Commun. 2018;9(1). doi:10.1038/s41467-018-04497-x

58. Cacciatore M, Magnani FG, Barbadoro F, et al. Thalamus and consciousness: a systematic review on thalamic nuclei associated with consciousness. Front Neurol. 2025;16. doi:10.3389/fneur.2025.1509668

59. A. Berry S R, Quan, Abreu. The AASM Manual for the Scoring of Sleep and Associated Events: Rules, Terminology and Technical Specifications. 2.6 versio.; 2020.

60. Delorme A, Makeig S. EEGLAB: An open source toolbox for analysis of single-trial EEG dynamics including independent component analysis. Journal of Neuroscience Methods. 2004;134(1):9–21. doi:10.1016/j.jneumeth.2003.10.009

61. Tadel F, Baillet S, Mosher JC, Pantazis D, Leahy RM. Brainstorm: A user-friendly application for MEG/EEG analysis. Computational Intelligence and Neuroscience. Published online 2011. doi:10.1155/2011/879716

62. Somervail R, Cataldi J, Stephan AM, Siclari F, Iannetti GD. Dusk2Dawn: an EEGLAB plugin for automatic cleaning of whole-night sleep electroencephalogram using Artifact Subspace Reconstruction. SLEEP. 2023;46(12):zsad208. doi:10.1093/sleep/zsad208

63. Bailey NW, Biabani M, Hill AT, et al. Introducing RELAX: An automated pre-processing pipeline for cleaning EEG data - Part 1: Algorithm and application to oscillations. Clinical Neurophysiology. 2023;149:178–201. doi:10.1016/j.clinph.2023.01.017

64. Richards JE, Sanchez C, Phillips-Meek M, Xie W. A database of age-appropriate average MRI templates. NeuroImage. Published online 2016. doi:10.1016/j.neuroimage.2015.04.055

65. Tzourio-Mazoyer N, Landeau B, Papathanassiou D, et al. Automated anatomical labeling of activations in SPM using a macroscopic anatomical parcellation of the MNI MRI single-subject brain. NeuroImage. Published online 2002. doi:10.1006/nimg.2001.0978

66. Pascual-Marqui RD. Standardized low-resolution brain electromagnetic tomography (sLORETA): Technical details. Methods and Findings in Experimental and Clinical Pharmacology. 2002;24(SUPPL. D):5–12.

67. Visalli A, Montefinese M, Viviani G, Finos L, Vallesi A, Ambrosini E. lmeEEG: Mass linear mixed-effects modeling of EEG data with crossed random effects. Journal of Neuroscience Methods. 2024;401:109991. doi:10.1016/j.jneumeth.2023.109991

68. Freedman D, Lane D. A Nonstochastic Interpretation of Reported Significance Levels. Journal of Business & Economic Statistics. 1983;1(4):292–298. doi:10.1080/07350015.1983.10509354

69. Spisák T, Spisák Z, Zunhammer M, et al. Probabilistic TFCE: A generalized combination of cluster size and voxel intensity to increase statistical power. NeuroImage. 2019;185:12–26. doi:10.1016/j.neuroimage.2018.09.078

