## Supplementary Table 6 for "Shared local brain dynamics in pediatric and adult NREM parasomnias"

| BRAIN LOBE | SPECIFIC BRAIN REGION | PREONSET | | POSTONSET | | AUTHOR/DATE | TECHNIQUE |
|---|---|---|---|---|---|---|---|
| FRONTAL | **Dorsolateral Prefrontal Cortex (Superior and Middle frontal gyri)** | | ↑ (low) delta | | ↓ (low) delta, ↑ 4-30 Hz, no HSDA | Flamand et al., 2018 | sEEG |
| | **Superior Frontal Gyrus** | | | | Slow waves (left) | Terzaghi et al., 2009 | sEEG |
| | **Middle Frontal Gyrus** | | | | Slow waves (right) | Terzaghi et al., 2012 | sEEG |
| | **Inferior Frontal Gyrus** | | | | Slow waves (left) | Terzaghi et al., 2009 | sEEG |
| | | | | | HSDA | Flamand et al., 2018 | sEEG |
| | **Frontal associative cortex** | | | | Regional cerebral blood flow < wake | Bassetti et al., 2000 | SPECT |
| | **Orbitofrontal Cortex** | | | | ↑ 4–10 Hz, HSDA | Flamand et al., 2018 | sEEG |
| | **Gyrus Rectus** | | | | No HSDA | Flamand et al., 2018 | sEEG |
| | **Dorsolateral Premotor Cortex** | | ↑ delta | | ↓ (low) delta, ↑ 5-27 Hz,no HSDA | Flamand et al., 2018 | sEEG |
| | **Supplementary Motor Area** | | ↑ delta | | ↓ (low) delta, no HSDA | Flamand et al., 2018 | sEEG |
| | | | | | Fast activity (right) | Terzaghi et al., 2012 | sEEG |
| | **Precentral Gyrus** | | | | Fast activity ~25 Hz (left) | Terzaghi et al., 2009 | sEEG |
| | | | ↑ (low) delta | | ↓ (low) delta, ↑ 17-30 Hz, no HSDA | Flamand et al., 2018 | sEEG |
| | **Precentral Operculum** | | ↑ delta | | No HSDA | Flamand et al., 2018 | sEEG |
| | **Anterior Cingulate Cortex** | | ↑ delta | | ↓ (low) delta, HSDA, no high-frequency | Flamand et al., 2018 | sEEG |
| | | | | | Fast activity (right) | Terzaghi et al., 2012 | sEEG |
| | **Midcingulate Cortex** | | ↑ delta (left) | | Fast activity (~25 Hz) (left) | Terzaghi et al., 2009 | sEEG |
| | | | | | ↓ (low) delta, no HSDA | Flamand et al., 2018 | sEEG |
| PARIETAL | **Posterior Cingulate Cortex** | | | | ↓ (low) delta, occasional HSDA | Flamand et al., 2018 | sEEG |
| | | | | | ↑ blood flow (compared to SWS) | Bassetti et al., 2000 | SPECT |
| | **Superior Parietal Lobule** | | | | Slow waves (left) | Terzaghi et al., 2009 | sEEG |
| | | | | | ↓ (low) delta, ↑ 7-12 Hz, HSDA | Flamand et al., 2018 | sEEG |
| | **Inferior Parietal Lobule** | | | | Slow waves (left) | Terzaghi et al., 2009 | sEEG |
| | | | | | ↓ (low) delta, HSDA | Flamand et al., 2018 | sEEG |
| | | | | | ↑ delta | Sarasso et al., 2014 | sEEG |
| | **Parietal associative cortex** | | | | ↓ blood flow (compared to wake) | Bassetti et al., 2000 | SPECT |
| | **Precuneus** | | | | ↓ (low) delta, ↑ 4-7 Hz, HSDA | Flamand et al., 2018 | sEEG |

| | | | | | | | |
|---|---|---|---|---|---|---|---|
| | **Postcentral Gyrus** | | | | ↓ (low) delta, ↑ 13-30 Hz, occasional HSDA | Flamand et al., 2018 | sEEG |
| | **Supramarginal Gyrus** | | | | HDSA | Flamand et al., 2018 | sEEG |
| | **Postcentral Operculum** | | ↑ (low) delta | | ↓ (low) delta, no HDSA | Flamand et al., 2018 | sEEG |
| **TEMPORAL** | **Amygdala** | | No changes | | Fast activity (right) | Terzaghi et al., 2012 | sEEG |
| | | | | | No HDSA | Flamand et al., 2018 | sEEG |
| | **Temporal Pole** | | Slow wave bursts | | Fast activity | Terzaghi et al., 2012 | sEEG |
| | **Superior Temporal Gyrus** | | ↑ delta | | ↓ (low) delta, ↑ 13-30 Hz, rare HSDA | Flamand et al., 2018 | sEEG |
| | **Middle Temporal Gyrus** | | ↑ delta | | ↓ (low) delta, ↑ 4-7 Hz, occasional HSDA | Flamand et al., 2018 | sEEG |
| | **Inferior Temporal Gyrus** | | ↑ delta | | ↓ (low) delta, no HSDA | Flamand et al., 2018 | sEEG |
| | **Hippocampus** | | ↑ (low) delta | | ↓ (low) delta, HSDA (50% cases), spindles (25% cases) | Flamand et al., 2018 | sEEG |
| | | | | | Spindles (right) | Terzaghi et al., 2012 | sEEG |
| | **Insula** | | ↑ (low) delta | | ↓ (low) delta, no HSDA | Flamand et al., 2018 | sEEG |
| | | | | | Fast activity (right) | Terzaghi et al., 2012 | sEEG |
| **OCCIPITAL** | **Occipital Cortex** | | No changes | | No changes, no HSDA | Flamand et al., 2018 | sEEG |
| **THALAMUS** | **Ventral Intermediate Nucleus** | | | | ↑ Beta, ↓Delta | Sarasso et al., 2014 | sEEG |
| | **Pulvinar** | | | | Occasional delayed HSDA | Flamand et al., 2018 | sEEG |
| **CEREBELLUM** | **Anterior Cerebellum** | | | | ↑ blood flow (> 25% compared to SWS) | Bassetti et al., 2000 | SPECT |

**Supplementary Table 6: Summary Of Stereo-EEG And Spect Findings By Brain Region**

Brain lobe – major brain lobe involved (e.g., frontal, temporal, parietal, occipital), specific brain region – anatomical sub-region within the lobe (e.g., dorsolateral prefrontal cortex, superior frontal gyrus), preonset – findings recorded before DoA motor onset, postonset – findings recorded after DoA onset, author/date – citation of the study reporting the findings, technique – method used to obtain the data (seeg = stereo-electroencephalography, spect = single-photon emission computed tomography). Blue = sleep-like activity; Red = wake-like activity; Violet = mixed activity
