## Supplementary Table 1 for "Shared local brain dynamics in pediatric and adult NREM parasomnias"

| ID | Sex | Age | DoA Subtype | Family history | Age at Onset | Frequency | Medications | Sleep comorbidities | Other major comorbidities |
|---|---|---|---|---|---|---|---|---|---|
| **A1** | M | 34.2 | CA | Yes | 4 | >1/week | None | Mild chronic sleep deprivation | None |
| **A2** | M | 28.5 | SW | No | 14 | >1/week | None | Mild OSA | None |
| **A3** | M | 36.0 | SW, SRSD | No | 32 (*) | 1/month | None | Sleeptalking, bruxism | GERD, Crohn's disease |
| **A4** | M | 27.3 | SW ST, SRSD | Yes | 2 | >1/week | None | Sleeptalking, hypnagogic hallucinations | None |
| **A5** | F | 31.8 | SW, ST | Yes | 10 | >1/month | None | Sleeptalking, hypnagogic hallucinations | None |
| **A6** | F | 27.0 | SW, ST | Yes | 8 | 1/month | Estro-Progestinic | Sleeptalking, hypnagogic hallucinations | Headache |
| **A7** | M | 39.8 | SW | Unknown | 6 | >1/week | Bisoprolol 2.5 mg (**) | Moderate OSA, sleeptalking, hypnagogic hallucinations | Hypertension |
| **A8** | F | 36.8 | CA | Yes | 6 | >1/year | None | None | None |
| **A9** | M | 21.6 | SW, CA | No | 5 | clusters | None | None | Migraine |
| **A10** | F | 28.4 | SW, ST | Yes | 6 | >1/week | L-Thyroxine 100 mcg, Estro-Progestinic | None | Hypotyroidism (thyroid agenesis) |
| **A11** | F | 29.9 | SW | Yes | 4 | >1/week | Zolpidem 2.5 mg as needed -on average 1/month (***) | Mild insomnia | None |
| **A12** | F | 25.4 | CA | Yes | 6 | >1/week | Estro-Progestinic | Mild insomnia | None |
| **A13** | M | 36.5 | SW | Yes | 20 (*) | >1/month | None | Bruxism | None |
| **A14** | F | 31.8 | SW | Yes | 7 | 1/month | None | None | GERD |
| **A15** | F | 29.3 | ST | Yes | 9 | >1/week | Estro-Progestinic | None | Migraine |
| **A16** | F | 23.0 | CA | Unknown | 8 | clusters | L-Thyroxine 125 mcg, Estro-Progestinic | None | Hypotyroidism, |
| **A17** | M | 37.7 | CA | Yes | 6 | >1/week | L-Thyroxine 150/175 mcg | Bruxism, mild OSA | Hypotyroidism |
| **A18** | M | 36.3 | SW, ST | No | 6 | 1/month | None | Moderate OSA | None |
| **A19** | F | 25.0 | SW, CA | Yes | 8 | everyday | None | None | None |
| **A20** | M | 34.1 | CA | No | 14 | >1/month | None | Mild OSA | None |
| **A21** | F | 23.6 | SW | No | ped | >1/week | Erenumab 70 mg (**) | None | Migraine |
| **A22** | F | 32.7 | SW, ST, CA | Yes | ped | >1/week | None | Mild insomnia | None |

**Supplementary Table 1: Clinical and Demographic Characteristics of Adult DoA Participants**

ID – Participant code; Sex – M = Male, F = Female; Age – Age at the time of enrollment (years); DoA: Disorders of Arousal, Subtype – CA = Confusional arousals, SW = Sleepwalking, ST = Sleep terrors, SRSD = Sleep-related sexual disorder; Family history – Presence of the disorder in family (Yes, No, Unknown); Age at onset – Age when symptoms began (years, or "ped" = pediatric onset); Frequency – Episode occurrence rate; Medications – Current treatments at the time of enrollment; Sleep comorbidities – Other diagnosed sleep disorders (OSA = obstructive sleep apnea); Other major comorbidities – Other physical health conditions (GERD = gastroesophageal reflux disease). (*) Uncertain episodes during childhood; (**) Medication withheld on the day of recording; discontinued three days prior; (***) One-month washout period. Note: A7 and A18 were excluded due to the finding of moderate OSA.
