## Supplementary Table 2 for "Shared local brain dynamics in pediatric and adult NREM parasomnias"

| ID | Sex | Age | DoA Subtype | Family history | Age at onset | Frequency | Medications | Sleep comorbidities | Other major comorbidities |
|---|---|---|---|---|---|---|---|---|---|
| **C1** | M | 13.3 | SW | Yes | 7 | 1/week | Melatonin 1 mg (*) | Mild insomnia | Separation anxiety |
| **C2** | M | 13.4 | SW, NT | Yes | 4 | >1/week | None | Sleep hyperhydrosis | None |
| **C3** | M | 11.2 | SW | Yes | 4 | >1/month | Salbutamol spray as needed (*) | None | Orchiopexy, amelanogenesis imperfecta, separation anxiety |
| **C4** | M | 15.4 | CA | Yes | 4 | >1/month | None | Sleeptalking, bruxism, mild insomnia | Talaxemia minor, tics, needle phobia |
| **C5** | M | 12.9 | SW, CA, NT | Yes | 3 | >1/week | None | None | None |
| **C6** | F | 7.3 | SW | No | 7 | Everyday | None | None | None |
| **C7** | F | 12.2 | SW | Yes | 3 | >1/week | None | Mild insomnia, sleeptalking, sleep hallucinations | Asthma, mild compulsive symptoms |
| **C8** | F | 7.2 | SW | Yes | 5 | >1/week | None | None | Asthma, epileptic abnormalities |
| **C9** | F | 8.3 | NT, SW | Yes | 5 | >1/month | None | None | None |
| **C10** | M | 5.5 | NT, CA | Yes | 6 | Everyday | None | None | None |
| **C11** | M | 8.6 | SW, CA | No | 7 | >1/week | None | Severe OSA, bruxism, bleeptalking | None |
| **C12** | M | 11.0 | SW, NT | Yes | 7 | Everyday | None | Flow-limitation | None |
| **C13** | M | 9.5 | SW, CA | No | 4 | >1/week | Laxatives as needed | Mild insomnia, sleep hyperhydrosis | Patent foramen ovale, autism spectrum disorder – level 1 |
| **C14** | M | 12.7 | SW | No | 9 | >1/week | None | Bruxism | Primary exercise headache |
| **C15** | M | 12.5 | SW | Yes | 6 | Everyday | Metylphenidate 10 mg (*) | Sleep hyperhydrosis | ADHD, Rolandic epilepsy (past) |
| **C16** | M | 8.6 | NT, SW | No | 5 | >1/week | None | None | None |
| **C17** | M | 12.3 | NT, CA, SW | No | 4 | Everyday | L-OH-Tryptophane 50 mg (*) | None | None |
| **C18** | F | 8.8 | SW | Yes | 6 | >1/month | None | Mild insomnia, flow-limitation | None |
| **C19** | F | 16.7 | NT, CA, SW | No | 6 | >1/week | Isotretinoinum 20 mg (*) | None | Acne |

**Supplementary Table 2: Clinical and Demographic Characteristics of Children DoA Participants**
ID – Participant code; Sex – M = Male, F = Female; Age – Age at the time of enrollment (years); DoA: Disorders of Arousal, Subtype – CA = Confusional arousals, SW = Sleepwalking, ST = Sleep terrors, Family history – Presence of the disorder in family (Yes, No, Unknown); Age at onset – Age when symptoms began (years, or "ped" = pediatric onset); Frequency – Episode occurrence rate; Medications – Current treatments at the time of enrollment; Sleep comorbidities – Other diagnosed sleep disorders (OSA = obstructive sleep apnea); Other major comorbidities – Other physical health conditions or neuro-psychiatric comorbidities – (ADHD = attention deficit hyperactivity disorder). (*) Medication withheld on the day of recording; discontinued three days prior. Note: C11 was excluded after the recording due to severe OSA for pediatric age and C8 due to the detection of isolated frontal sharp waves, more prominent on the right side, visible during wake and all sleep stages, without clinical correlates. C12 and C18 exhibited flow-limitation but had an AHI <3 and were therefore retained for analysis.
