## Supplementary Table 3 for "Shared local brain dynamics in pediatric and adult NREM parasomnias"

| ID | LMSI (n/h) | PLMSI (n/h) | AHI (n/h) | TIB (min) | TST (min) | SL (min) | REML (min) | WASO (min) | SE (%) | N1 (min) | N1 (%) | N2 (min) | N2 (%) | N3 (min) | N3 (%) | REM (min) | REM (%) | AI (n/h) | NREM-AI (n/h) | REM-AI (n/h) |
|---|---|---|---|---|---|---|---|---|---|---|---|---|---|---|---|---|---|---|---|---|
| **A1** | 6.0 | 0.0 | 3.6 | 242.1 | 224.1 | 4.0 | 99.0 | 14.0 | 92.5 | 15.1 | 6.7 | 130.5 | 58.2 | 58.0 | 25.9 | 20.5 | 9.2 | 8.8 | 14.7 | 11.7 |
| **A2** | 18.4 | 4.8 | 7.5 | 464.0 | 411.6 | 9.5 | 180.5 | 42.9 | 88.7 | 47.5 | 11.5 | 255.6 | 62.1 | 28.9 | 7.0 | 79.6 | 19.3 | 16.5 | 29.6 | 12.1 |
| **A3** | 24.2 | 3.7 | 8.9 | 486.5 | 382.0 | 14.0 | 97.5 | 26.5 | 78.5 | 27.5 | 7.2 | 162.0 | 42.4 | 73.5 | 19.2 | 119.0 | 31.1 | 15.6 | 31.9 | 8.6 |
| **A4** | 31.9 | 21.0 | 5.1 | 405.0 | 373.3 | 8.5 | 144.5 | 23.2 | 92.2 | 47.2 | 12.6 | 171.0 | 45.8 | 64.7 | 17.3 | 90.5 | 24.2 | 15.8 | 29.3 | 8.6 |
| **A5** | 27.8 | 11.4 | 0.1 | 443.0 | 387.7 | 6.0 | 143.5 | 49.4 | 87.5 | 43.8 | 11.3 | 200.5 | 51.7 | 59.5 | 15.3 | 83.8 | 21.6 | 22.3 | 33.6 | 23.6 |
| **A6** | 7.2 | 0.6 | 0.5 | 436.3 | 377.8 | 8.8 | 126.2 | 49.6 | 86.6 | 29.5 | 7.8 | 201.2 | 53.2 | 82.2 | 21.8 | 65.0 | 17.2 | 14.3 | 26.1 | 11.1 |
| **A7** | 9.8 | 0.7 | 23.5 | 417.6 | 332.4 | 3.5 | 93.0 | 81.7 | 79.6 | 38.1 | 11.5 | 125.9 | 37.9 | 74.0 | 22.3 | 94.4 | 28.4 | 18.9 | 36.3 | 5.7 |
| **A8** | 14.4 | 2.3 | 1.2 | 436.0 | 363.0 | 5.0 | 137.5 | 73.0 | 83.3 | 18.0 | 5.0 | 145.5 | 40.1 | 115.5 | 31.8 | 84.0 | 23.1 | 18.8 | 43.4 | 5.7 |
| **A9** | 10.7 | 2.0 | 2.2 | 443.4 | 403.4 | 9.0 | 85.5 | 28.5 | 91.0 | 15.4 | 3.8 | 138.0 | 34.2 | 157.5 | 39.0 | 92.5 | 22.9 | 15.5 | 26.2 | 13.6 |
| **A10** | 16.8 | 6.4 | 1.3 | 401.4 | 358.4 | 10.0 | 196.0 | 33.0 | 89.3 | 35.1 | 9.8 | 165.4 | 46.2 | 72.9 | 20.3 | 84.9 | 23.7 | 19.6 | 36.0 | 13.4 |
| **A11** | 20.5 | 12.5 | 0.0 | 408.0 | 361.6 | 2.0 | 87.0 | 44.4 | 88.6 | 19.9 | 5.5 | 159.5 | 44.1 | 79.6 | 22.0 | 102.6 | 28.4 | 11.3 | 12.1 | 13.4 |
| **A12** | 17.0 | 2.5 | 0.0 | 397.4 | 228.2 | 19.5 | 232.5 | 149.8 | 57.4 | 46.0 | 20.2 | 79.7 | 34.9 | 66.0 | 28.9 | 36.5 | 16.0 | 23.9 | 40.1 | 8.2 |
| **A13** | 7.2 | 1.2 | 2.6 | 388.0 | 342.5 | 8.0 | 106.0 | 37.5 | 88.3 | 25.0 | 7.3 | 135.5 | 39.6 | 90.5 | 26.4 | 91.5 | 26.7 | 13.5 | 30.6 | 5.9 |
| **A14** | 11.9 | 4.4 | 0.0 | 371.0 | 268.8 | 3.5 | 114.5 | 98.7 | 72.5 | 17.7 | 6.6 | 151.1 | 56.2 | 52.9 | 19.7 | 47.1 | 17.5 | 17.9 | 29.8 | 19.1 |
| **A15** | 15.5 | 1.4 | 0.0 | 404.6 | 324.1 | 9.5 | 152.5 | 71.0 | 80.1 | 51.1 | 15.8 | 178.0 | 54.9 | 21.0 | 6.5 | 74.0 | 22.8 | 23.9 | 44.1 | 16.2 |
| **A16** | 15.3 | 4.7 | 3.3 | 374.7 | 340.6 | 12.5 | 162.0 | 21.6 | 90.9 | 13.3 | 3.9 | 165.9 | 48.7 | 87.8 | 25.8 | 73.5 | 21.6 | 16.6 | 23.4 | 25.3 |
| **A17** | 10.4 | 2.4 | 6.9 | 458.2 | 417.8 | 9.5 | 47.0 | 30.8 | 91.2 | 41.0 | 9.8 | 191.7 | 45.9 | 72.0 | 17.2 | 113.2 | 27.1 | 23.1 | 50.0 | 11.1 |
| **A18** | 31.4 | 16.8 | 24.2 | 470.2 | 379.4 | 19.5 | 102.5 | 71.3 | 80.7 | 30.5 | 8.0 | 186.7 | 49.2 | 81.8 | 21.6 | 80.4 | 21.2 | 28.5 | 63.8 | 3.7 |
| **A19** | 6.0 | 0.0 | 0.5 | 389.4 | 334.9 | 29.0 | 98.5 | 25.5 | 86.0 | 23.7 | 7.1 | 190.7 | 56.9 | 67.3 | 20.1 | 53.2 | 15.9 | 13.6 | 25.1 | 4.5 |
| **A20** | 10.3 | 0.7 | 9.0 | 462.8 | 377.9 | 25.5 | 94.0 | 59.4 | 81.7 | 25.7 | 6.8 | 178.9 | 47.3 | 79.0 | 20.9 | 94.3 | 25.0 | 16.4 | 25.4 | 18.4 |
| **A21** | 18.7 | 2.5 | 0.2 | 498.0 | 338.0 | 20.5(*) | 77.0 | 125.0 | 67.9 | 43.3 | 12.8 | 178.7 | 52.9 | 64.5 | 19.1 | 51.5 | 15.2 | 21.8 | 41.0 | 4.7 |
| **A22** | 5.1 | 0.0 | 0.0 | 369.7 | 215.2 | 16.0 | 386.5 | 138.5 | 58.2 | 54.0 | 25.1 | 98.7 | 45.9 | 14.0 | 6.5 | 48.5 | 22.5 | 56.3 | 129.5 | 5.0 |

**Supplementary Table 3: Sleep Study Summary Table of Adult DoA Participants.**
ID = Subject identifier; LMSI (n/h) = Limb Movement Sleep Index (number/hour); PLMSI (n/h) = Periodic Limb Movement Sleep Index (number/hour); AHI (n/h) = Apnea–Hypopnea Index (number/hour); TIB (min) = Time in Bed; TST (min) = Total Sleep Time; SL (min) = Sleep Latency; REML (min) = REM Latency; WASO (min) = Wake After Sleep Onset; SE (%) = Sleep Efficiency; N1, N2, N3, REM (min) = minutes spent in each sleep stage; N1, N2, N3, REM (%) = percentage of TST in each sleep stage; AI (n/h) = Arousal Index; NREM AI (n/h) = NREM Arousal Index; REM AI (n/h) = REM Arousal Index.
(*) – Sleep latency was underestimated due to technical issues occurring between light-off and sleep onset.
