## Supplementary Table 4 for "Shared local brain dynamics in pediatric and adult NREM parasomnias"

| ID | LMSI (n/h) | PLMSI (n/h) | AHI (n/h) | TIB (min) | TST (min) | SL (min) | REML (min) | WASO (min) | SE (%) | N1 (min) | N1 (%) | N2 (min) | N2 (%) | N3 (min) | N3 (%) | REM (min) | REM (%) | AI (n/h) | NREM AI (n/h) | REM AI (n/h) |
|---|---|---|---|---|---|---|---|---|---|---|---|---|---|---|---|---|---|---|---|---|
| **C1** | 9.8 | 4.0 | 1.9 | 418.4 | 355.5 | 4.5 | 142.0 | 58.4 | 85.0 | 32.0 | 9.0 | 117.3 | 33.0 | 147.4 | 41.5 | 58.8 | 16.5 | 16.2 | 29.5 | 11.2 |
| **C2** | 11.8 | 0.6 | 0.3 | 448.6 | 421.0 | 8.0 | 141.0 | 19.6 | 93.9 | 20.0 | 4.8 | 160.5 | 38.1 | 149.5 | 35.5 | 91.0 | 21.6 | 10.1 | 12.7 | 16.5 |
| **C3** | 16.8 | 4.5 | 0.3 | 483.4 | 391.4 | 62.0 | 227.5 | 30.0 | 81.0 | 20.6 | 5.3 | 170.6 | 43.6 | 135.4 | 34.6 | 64.9 | 16.6 | 11.3 | 22.8 | 4.6 |
| **C4** | 15.8 | 3.2 | 1.6 | 445.4 | 417.8 | 6.7 | 121.8 | 20.9 | 93.8 | 32.5 | 7.8 | 148.5 | 35.5 | 151.8 | 36.3 | 85.0 | 20.3 | 14.2 | 26.3 | 9.2 |
| **C5** | 15.0 | 6.0 | 0.2 | 491.2 | 348.0 | 10.0 | 244.0 | 142.2 | 70.8 | 22.6 | 6.5 | 152.4 | 43.8 | 104.0 | 29.9 | 69.0 | 19.8 | 12.8 | 24.5 | 2.6 |
| **C6** | 14.4 | 3.3 | 1.3 | 543.2 | 493.9 | 18.5 | 103.5 | 30.8 | 90.9 | 39.0 | 7.9 | 169.2 | 34.3 | 166.0 | 33.6 | 119.7 | 24.2 | 10.6 | 17.6 | 12.0 |
| **C7** | 14.7 | 2.1 | 0.6 | 507.0 | 419.0 | 41.5 | 78.0 | 46.5 | 82.6 | 29.5 | 7.0 | 141.0 | 33.7 | 132.0 | 31.5 | 116.5 | 27.8 | 14.6 | 19.8 | 17.0 |
| **C8** | 22.4 | 9.8 | 2.1 | 517.4 | 458.9 | 30.5 | 169.5 | 28.0 | 88.7 | 25.2 | 5.5 | 172.3 | 37.5 | 139.2 | 30.3 | 122.3 | 26.6 | 12.8 | 25.0 | 7.9 |
| **C9** | 17.8 | 10.0 | 0.6 | 428.0 | 379.3 | 7.7 | 85.5 | 41.0 | 88.6 | 37.6 | 9.9 | 157.5 | 41.5 | 115.2 | 30.4 | 69.0 | 18.2 | 11.1 | 13.1 | 12.2 |
| **C10** | 22.8 | 4.8 | 0.2 | 434.5 | 373.0 | (*) | 137.5 | 61.5 | 85.8 | 26.0 | 7.0 | 124.0 | 33.2 | 143.5 | 38.5 | 79.5 | 21.3 | 11.7 | 18.0 | 13.6 |
| **C11** | 25.9 | 4.9 | 13.8 | 461.1 | 388.6 | 11.5 | 135.0 | 61.0 | 84.3 | 39.1 | 10.1 | 146.6 | 37.7 | 115.0 | 29.6 | 88.0 | 22.6 | 13.1 | 23.9 | 7.5 |
| **C12** | 35.9 | 19.4 | 1.9 | 465.9 | 403.5 | 9.5 | 152.0 | 52.9 | 86.6 | 35.7 | 8.8 | 121.0 | 30.0 | 141.3 | 35.0 | 105.5 | 26.1 | 19.3 | 34.2 | 25.6 |
| **C13** | 11.2 | 2.6 | 2.5 | 499.6 | 438.0 | 39.5 | 72.5 | 22.1 | 87.7 | 39.4 | 9.0 | 163.3 | 37.3 | 129.5 | 29.6 | 105.9 | 24.2 | 11.5 | 24.9 | 7.4 |
| **C14** | 8.4 | 0.0 | 0.4 | 476.4 | 404.4 | 57.0 | 153.0 | 15.0 | 84.9 | 13.9 | 3.4 | 195.0 | 48.2 | 98.5 | 24.4 | 97.0 | 24.0 | 14.2 | 21.1 | 17.9 |
| **C15** | 9.0 | 0.9 | 0.6 | 474.3 | 420.7 | 37.0 | 196.0 | 16.5 | 88.7 | 12.2 | 2.9 | 185.9 | 44.2 | 150.8 | 35.9 | 71.8 | 17.1 | 9.6 | 13.4 | 16.7 |
| **C16** | 7.0 | 0.0 | 2.2 | 481.7 | 437.0 | 3.5 | 123.5 | 41.2 | 90.7 | 37.7 | 8.6 | 154.0 | 35.2 | 120.3 | 27.5 | 125.0 | 28.6 | 14.8 | 29.2 | 12.5 |
| **C17** | 12.1 | 4.9 | 0.8 | 487.9 | 429.5 | 15.0 | 44.5 | 21.9 | 88.0 | 18.0 | 4.2 | 193.6 | 45.1 | 118.5 | 27.6 | 99.4 | 23.1 | 10.6 | 16.7 | 9.1 |
| **C18** | 5.8 | 0.0 | 0.0 | 412.9 | 397.0 | 4.5 | 229.0 | 11.4 | 96.1 | 7.6 | 1.9 | 195.0 | 49.1 | 116.6 | 29.4 | 77.8 | 19.6 | 6.3 | 9.4 | 8.5 |
| **C19** | 1.7 | 0.0 | 0.1 | 478.3 | 431.8 | 4.0 | 75.5 | 37.5 | 90.3 | 22.5 | 5.2 | 182.3 | 42.2 | 118.5 | 27.4 | 108.5 | 25.1 | 10.8 | 23.4 | 6.1 |

**Supplementary Table 4: Sleep Study Summary Table of Children DoA Participants.**
ID = Subject identifier; LMSI (n/h) = Limb Movement Sleep Index (number/hour); PLMSI (n/h) = Periodic Limb Movement Sleep Index (number/hour); AHI (n/h) = Apnea–Hypopnea Index (number/hour); TIB (min) = Time in Bed; TST (min) = Total Sleep Time; SL (min) = Sleep Latency; REML (min) = REM Latency; WASO (min) = Wake After Sleep Onset; SE (%) = Sleep Efficiency; N1, N2, N3, REM (min) = minutes spent in each sleep stage; N1, N2, N3, REM (%) = percentage of TST in each sleep stage; AI (n/h) = Arousal Index; NREM AI (n/h) = NREM Arousal Index; REM AI (n/h) = REM Arousal Index.
(*) – Recording commenced after the participant had already fallen asleep, owing to technical issues.
