## Supplementary Table 5 for "Shared local brain dynamics in pediatric and adult NREM parasomnias"

| | | DOA episodes | Physiological Motor arousals |
|---|---|---|---|
| **Adult group** | | | |
| | Subjects with episodes (n) | 12 (60%) | 18 (90%) |
| | Number of episodes (n) | 34 | 68 |
| | Episodes per subject (mean ± SD, n) | 1.7 ± 2.0 | 3.4 ± 2.5 |
| | Duration (mean ± SD, s) | 16.0 ± 8.4 | 12.6 ± 10.1 |
| | Latency from onset (mean ± SD, min) | 115.7 ± 96.8 | 143.3 ± 105.3 |
| **Children group** | | | |
| | Subjects with episodes (n) | 15 (82%) | 16 (94%) |
| | Number of episodes (n) | 50 | 78 |
| | Episodes per subject (mean ± SD, n) | 2.9 ± 2.5 | 4.6 ± 2.9 |
| | Duration (mean ± SD, s) | 31.0 ± 16.2 | 14.8 ± 9.5 |
| | Latency from onset (mean ± SD, min) | 144.1 ± 87.1 | 136.2 ± 93.1 |

**Supplementary Table 5: DoA Episodes and Physiological Motor Arousals in Adult and Children Participants**
Comparison between DoA episodes and physiological motor arousals in adults and children. Values are reported as counts (n), percentages (%), or means ± standard deviations (SD). “Subjects with episodes” refers to participants presenting at least one recorded episode. “Number of episodes” is the total count observed. “Episodes per subject” represents the mean number of episodes per participant. “Duration” is the mean length of episodes in seconds. “Latency from onset” indicates the mean time from sleep onset to the first recorded episode, expressed in minutes.
